# Prioritization of semantic over visuo-perceptual aspects in multi-item working memory

**DOI:** 10.1101/2022.06.29.498168

**Authors:** Casper Kerrén, Juan Linde-Domingo, Bernhard Spitzer

## Abstract

Working Memory (WM) keeps information temporarily available for upcoming tasks. How the contents of WM are distinguished from perceptual representations on the one hand, and from long-term memories on the other, is still debated. Here, we leveraged recent evidence for a reversal of processing dynamics when retrieving episodic long-term memories as opposed to perceiving visual input. In two experiments (n=75 and n=103), we asked participants to hold one or more items in WM and to report their low-level perceptual and high-level semantic qualities. In both experiments, we found faster responses to the items’ semantic qualities, indicating prioritization over visuo-perceptual aspects, when two or more items were held concurrently in WM. These dynamics of accessing information in multi-item WM were akin to those in retrieving episodic long-term memories and opposite to those in processing visual inputs. Little to no semantic prioritization was evident during single-item maintenance, consistent with a strictly capacity-limited focus of attention within which WM information can be transformed into a prospective action plan.

## Introduction

Adaptive cognition critically hinges on information no longer available in the environment. The capacity to keep such information present “in one’s mind” and available for mental operations is often referred to as Working Memory (WM)^1,2^. A key question in understanding WM function concerns the nature of the retained information, and the ways in which it differs—or not—from what is encoded during perception^3–5^ or stored in long-term memory (LTM)^6–8^.

Neuroscientific work in humans and other animals has offered insights into WM processing during simple delay tasks, where often only a single stimulus or feature must be remembered^2,3,9^. Evidence has accumulated that WM information can be represented in the same brain circuits that are involved in sensory perception of this information^4,10–13^. In such a “sensory recruitment” framework of WM, working memories may share similarities with sensory percepts. However, there is also ample evidence that perceptual information is reformatted during WM storage^5,14–18^, for instance, into an abstraction of the task-relevant stimulus information^19^, and/or into a prospective response plan for the upcoming task^2,9,20^.

While many neuroscientific studies have focused on single-item maintenance, it is widely assumed that WM can maintain multiple pieces of information concurrently^21–23^, including information temporarily outside the focus of attention^6,24,25^. The nature of such “unattended” WM storage is intensely debated. Theoretical proposals include that unattended contents may temporarily persist in “activity-silent’’ short-term synaptic engrams^26,27^, or that they are redistributed across brain areas that may retain the information at different levels of abstraction^28^ (but see refs.^19,29^). Another, more long-standing view is that WM storage outside the focus of attention may rely on information in LTM^6,7,30^. Extending this idea, it has recently been proposed that unattended (or activity-silent) WM may recruit the same neurocognitive processes as episodic LTM^8^. Interestingly, however, it is not commonly assumed that LTM would contribute to the concurrent maintenance of multiple items (within the capacity limits of WM, e.g., Jeneson & Squire, 2012) between which attention must be shared.

Studies of LTM have shown that there are qualitative differences in how active mental representations are reconstructed from episodic memory, compared to when they emerge during perception^31,32^. During perception, the low-level visual features of a stimulus (e.g., its shape or color) are known to be processed earlier than its high-level conceptual aspects (e.g., its semantic category)^33,34,34–36^. Intriguingly, these temporal dynamics appear to be reversed in episodic LTM retrieval, where a memory’s semantic aspects tend to be retrieved faster than its visuo-perceptual details^31,32^ (see refs.^37,38^ for similar findings in visual imagery). These findings have been interpreted in support of the view that episodic LTM is inherently (re-)constructive^39–41^, possibly with the hippocampus triggering a processing cascade that follows the reverse trajectory of visual perception^31,42^. In contrast, little is known yet about the temporal dynamics of accessing information in WM, and whether they resemble those of visual perception, or those in episodic LTM retrieval.

Here, we took advantage of experimental techniques introduced in recent LTM work^31,32^ to examine whether visuo-perceptual or semantic information is prioritized when accessing visual object information in WM. A priori, the following three scenarios seem possible: if WM retains a concrete visual memory of the presented stimuli, we may expect faster access to their visuo-perceptual than to their semantic aspects. In contrast, if WM retains prospective action plans (i.e., anticipated responses in the upcoming test) we expect no prioritization of either perceptual or semantic dimensions to be detectable in behavior, since the response plans associated with the two dimensions would be in the same format (e.g., response contingencies^43^). We would thus expect no difference in their execution time when probed at test. Likewise, we would expect no differences in response time if the task-relevant stimulus aspects had been recoded into a verbal format, which we would expect to be equally accessible for either dimension. Critically, however, if WM storage relies on formats more akin to those in episodic LTM, we expect faster access to semantic than to visuo-perceptual aspects, as has previously been reported in episodic retrieval^31^.

We tested for these scenarios both in single-item maintenance under full attention and in maintenance of multiple items between which attention had to be shared. We found a clear prioritization of semantic aspects in multi-item maintenance, regardless of whether the items were probed with verbal (Experiment 1) or with spatial cues (Experiment 2). These findings indicate processing dynamics in WM that are more akin to those in episodic LTM retrieval than to those in visual perception. Semantic prioritization was less evident in single-item maintenance, in line with the notion of a single-item focus of attention within which WM information may take the form of prospective action plans.

## Results

### Experiment 1

Participants (n = 75) performed a WM task in which they were asked to associate visual stimuli with action verbs (Fig. 1b, *left*). The stimuli were either color photographs or line-drawings (perceptual dimension) of either animate or inanimate objects (semantic dimension; Fig. 1a). Either one, two, or three verb-stimulus pairs were presented (load 1-3, randomly varied across trials) until participants were prompted with one of the verbs (randomly selected) to report the attributes (perceptual and semantic) of the associated stimulus (Fig. 1b, *right*). Thus, load 1 trials allowed participants to report directly on the only stimulus in WM, whereas loads 2 and 3 required accessing the probed stimulus via the associated cue word (see Experiment 2 for results with a different, non-verbal cueing procedure).

**Figure 1.**
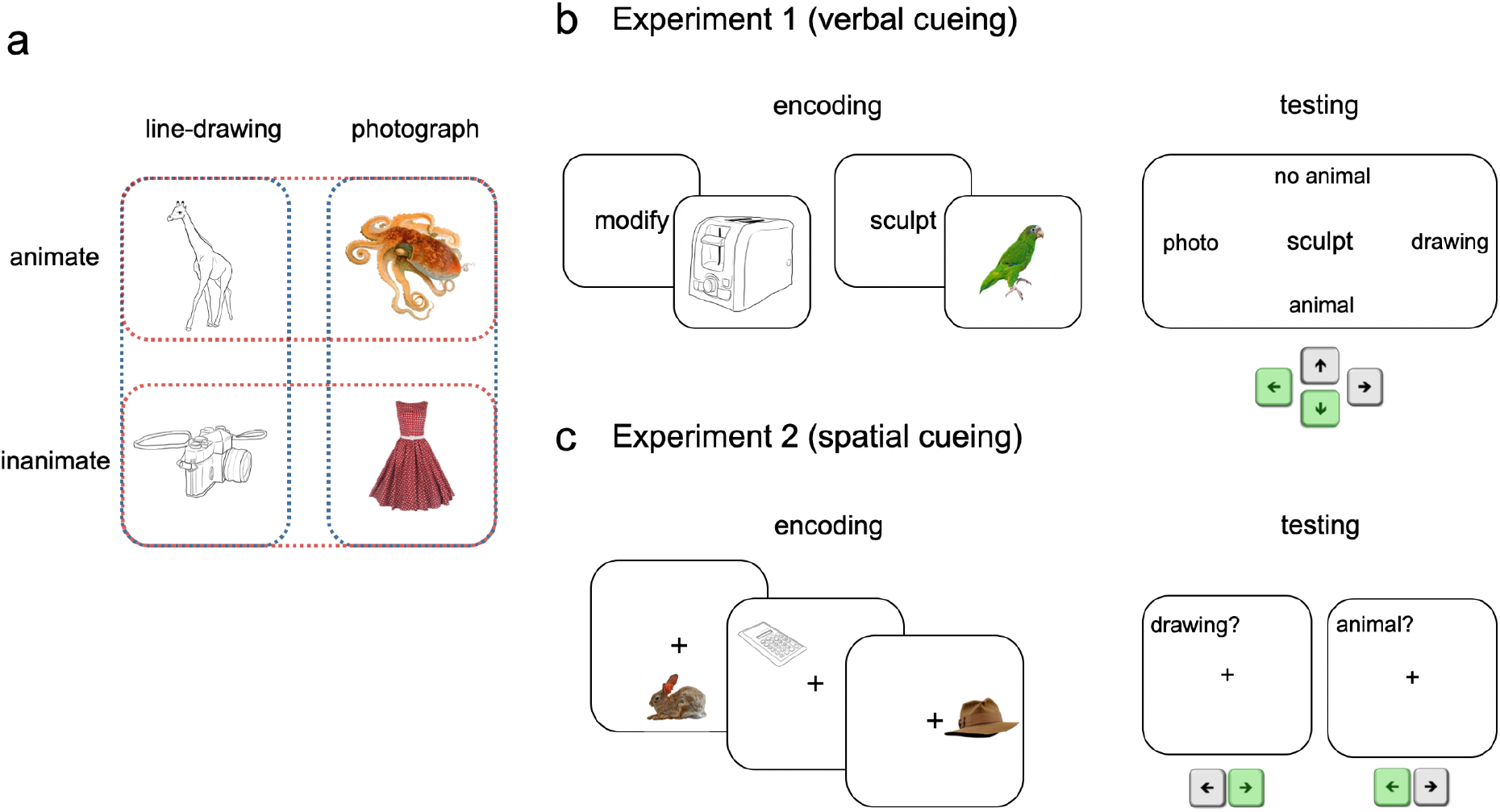
Stimuli and tasks. ***a***, Example stimuli. The to-be-remembered stimuli were line-drawings and color photographs (perceptual dimension, blue outlines) of animate and inanimate items (semantic dimension, red outlines). ***b***, Example trial in Experiment 1. At encoding (*left*), participants (n=75) were asked to associate each stimulus with an action verb. The number of verb-stimulus associations to be memorized on a trial was randomly varied (1-3 pairs). At test (*right*), one of the action verbs (here, “sculpt”) was displayed at center and participants were asked to indicate the perceptual and semantic aspects of the associated stimulus via key presses (two responses). ***c***, In Experiment 2, participants (n=103) were asked to memorize a series of stimuli in different locations (*left*). After 1-4 stimuli (randomly varied), participants were cued to indicate the perceptual and semantic aspects (serial order randomly varied) of the stimulus that had been presented in the cued location (here, the drawing of a calculator) with yes/no responses (*right*). In both experiments, participants were instructed to respond as fast and accurately as possible.

We first examined the overall speed (response times, RTs) and accuracy (percentages correct) with which participants reported the perceptual and the semantic aspects of the probed stimulus (Fig. 2a,d). On average over all conditions, the semantic aspect (animal or no animal) was reported faster than the perceptual aspect (photograph or drawing; Fig. 2a, M = 2.138 ms, SE = 40 ms vs. M = 2.268 ms, SE = 34 ms; Z = 4.76, r = .55, *p* = <.001; Wilcoxon signed-rank test). Furthermore, the semantic aspect was also reported more accurately than the perceptual aspect (Fig. 2d; M = 88.80%, SE = .79% compared to M = 84.74%, SE = .93%; Z = -4.79, r = -.55, *p* < .001, Wilcoxon signed-rank test). Together, here in a multi-item WM task, we observed a prioritization of the memory items’ semantic aspects over their perceptual aspects, which resembles previous findings in studies of episodic LTM retrieval^31,32^.

**Figure 2.**
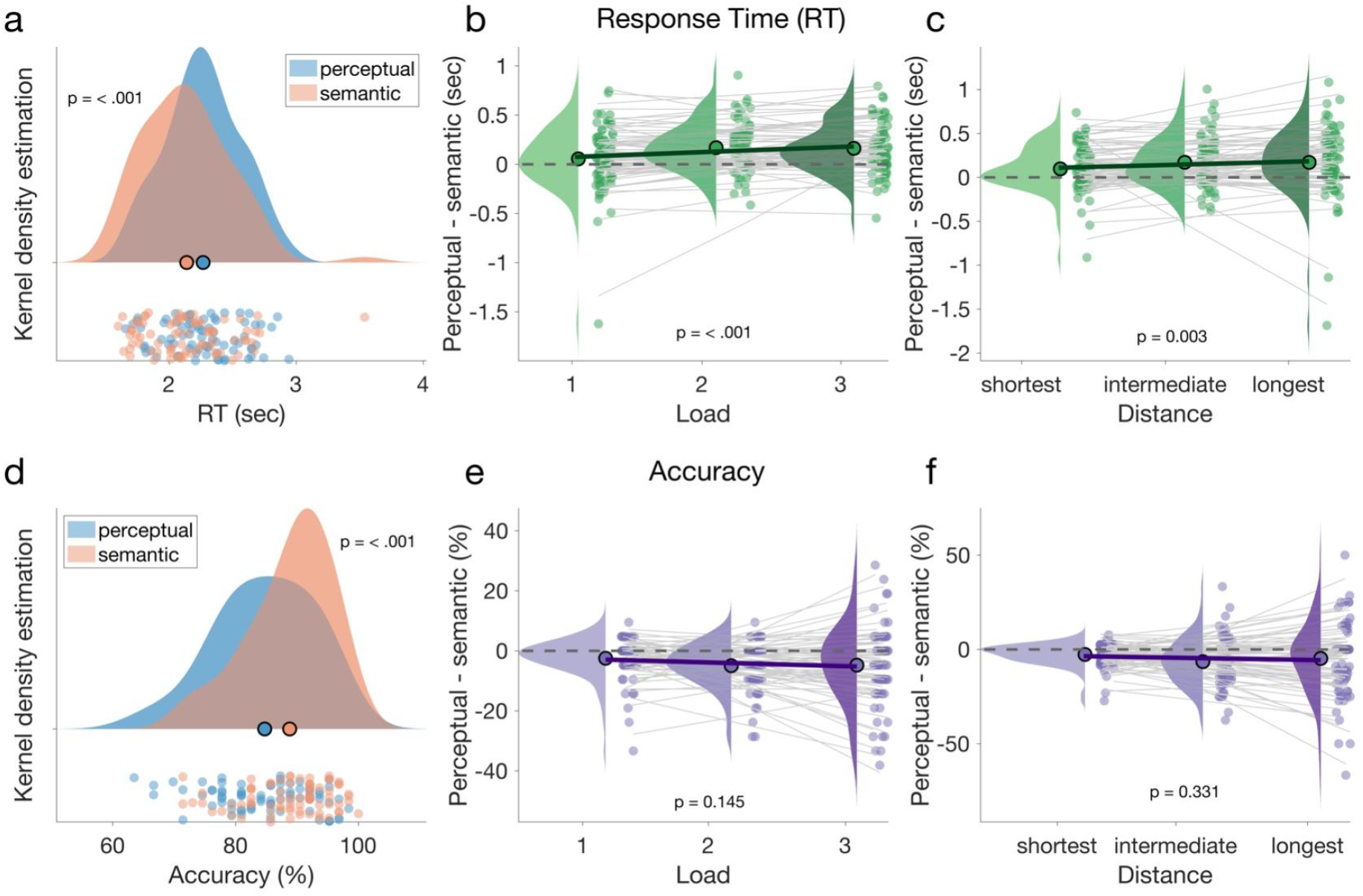
Response times and accuracy results, Experiment 1. **a**, Overall response times (RT), collapsed over load conditions (1-3), for the perceptual (photo/drawing; *blue*) and semantic (animal/no animal; *red*) aspects. Large dots: mean RT. Density plot illustrates RT distribution over participants. Small dots at the bottom show individual participant results. Participants on average reported the semantic aspect faster than the perceptual aspect. The p-value indicates significance of paired comparison. **b**, Difference in response times (y-axis; perceptual - semantic) across the three load conditions (x-axis). Positive values indicate relatively faster reports of the semantic aspect. Solid black line shows the average linear fit and thinner gray lines each participant’s linear fit. The p-value at the bottom indicates significance of the difference in linear slope from zero. **c**, Same as *b*, but grouping trials according to temporal distance between encoding and test (x-axis; see *Results* for details). Semantic prioritization increased both with WM load (*b*) and with temporal distance from test (*c*). **d-f**, Same as *a-c*, but for accuracy (proportion correct responses). The accuracy data overall mirrored the patterns in response times (*a*-*c*), however, the effects of load (*e*) and temporal distance (*f*) were not statistically significant.

Next, we asked to what extent the prioritization of semantic over visuo-perceptual information was modulated by WM load, that is, by the number of stimuli (1-3) that were to be maintained on a given trial. Indeed, a linear trend analysis of the RT differences (perceptual-semantic) showed that the prioritization of semantic aspects increased across loads 1-3 (Fig. 2b; load 1, M = 6ms, SE = 38ms; load 2, M = 169ms, SE = 28ms; load 3, M = 162ms, SE = 29ms; Z = 3.74, r = .43, *p* < .001; Wilcoxon signed-rank test of linear slope against zero). A similar trend in accuracy (in terms of a relative decrease in the accuracy of perceptual compared to semantic reports; Fig. 2e), was not statistically significant (load 1, M = -2.48%, SE = .76%; load 2, M = -4.95%, SE = .86%; load 3, M = -4.76%, SE = 1.47%, Z = -1.46,, r = -.17; *p* = .145). The RT effects observed in Exp. 1 could have been due to tendencies to report the semantic aspect first. To test for this, we examined the proportion of trials on which the semantic aspect (animate/inanimate) was reported before the visual aspect (photo/drawing) across load levels. While overall, participants indeed showed a tendency to report the semantic aspect first (mean = 59.30%, SE = 1.99%, Z = 4.13, r = .48, p < .001; Wilcoxon signed-rank test against 50%), we found no evidence for this tendency to increase across load levels (Z = .42, r = .05, p = .68; Wilcoxon signed-rank test of linear slope against zero). The load-dependent RT findings (Fig. 2b) can thus not be explained by differences in response order between the conditions of interest (see also Experiment 2 for results with a different test procedure).

An alternative way of interpreting the results of Exp. 1 is not in terms of the number of maintained items (i.e., load), but in terms of the delay between encoding and test, that is, the time during which a memory was temporarily unattended while processing other information. To this end, we grouped trials into “short”, “intermediate”, and “long” distance conditions. The short distance condition included all trials in which the most recently presented item was tested. Intermediate distances combined those 2- and 3-item trials in which the second-last presented item was tested, and the long-distance condition included those 3-item trials in which the first-presented item was tested. Note that the distances are moderately correlated with load levels (r=0.5). As expected, the semantic prioritization in RTs grew with the distance between encoding and test (Fig. 2c; short, M = 97ms, SE = 30ms; intermediate, M = 171ms, SE = 32ms; long, M = 170ms, SE = 46ms, Z = 3.03, r = .35 *p* = .0025). Again, there was no analogue significant effect in accuracy (Fig. 2f; short, M = -2.67%, SE = .67%; intermediate, M = -6.47%, SE = 1.44%; long, M = -4.77%, SE = 2.20%; Z = -.97, r = -.11, *p* = .3308).

In sum, Exp. 1 showed a prioritization of semantic aspects when accessing information in multi-item WM. However, a potential limitation of Exp. 1 was that the cueing with action verbs may have promoted associations that were predominantly semantic in nature. In our second experiment, we examined whether similar effects would also arise in a more conventional visual WM task setting where items are to be maintained in different spatial locations^44–47^.

### Experiment 2

The layout of Exp. 2 (n=103) was similar to that of Exp. 1, with the key difference being that the memory items were presented without action verbs at different locations on the screen (Fig. 1c). After presentation of 1-4 stimuli (the number randomly varied), participants were prompted to recall the perceptual and semantic aspects of the stimulus they remembered at the cued location (Fig 1c, *right*). The report was in the form of yes/no responses to randomized probe questions (see *Materials and Methods*) and we again focused on the speed (RTs) and accuracy (percentage correct) of participants’ responses.

The overall pattern of RTs and accuracy (Fig. 3) strongly resembled that in Exp. 1. On average in Exp. 2, participants were again faster and more accurate in reporting the memory items’ semantic qualities (animate/inanimate) as compared to their visual appearance (photo/drawing; Fig. 3a,d; RT: M = 1154ms, SE = 28ms and M = 1205ms, SE = 27ms; Z = 3.88, r = .38; *p* = .001, Wilcoxon signed-rank test; percentage correct: M = 89.35%, SE = .92% and M = 87.32%, SE = 1.03%; Z = -3.70, r = -.36, *p =* .002). Thus, we likewise found a prioritization of semantic aspects in a more conventional WM task, where a small number of stimuli (here 1-4) was to be remembered at their spatial location.

**Figure 3.**
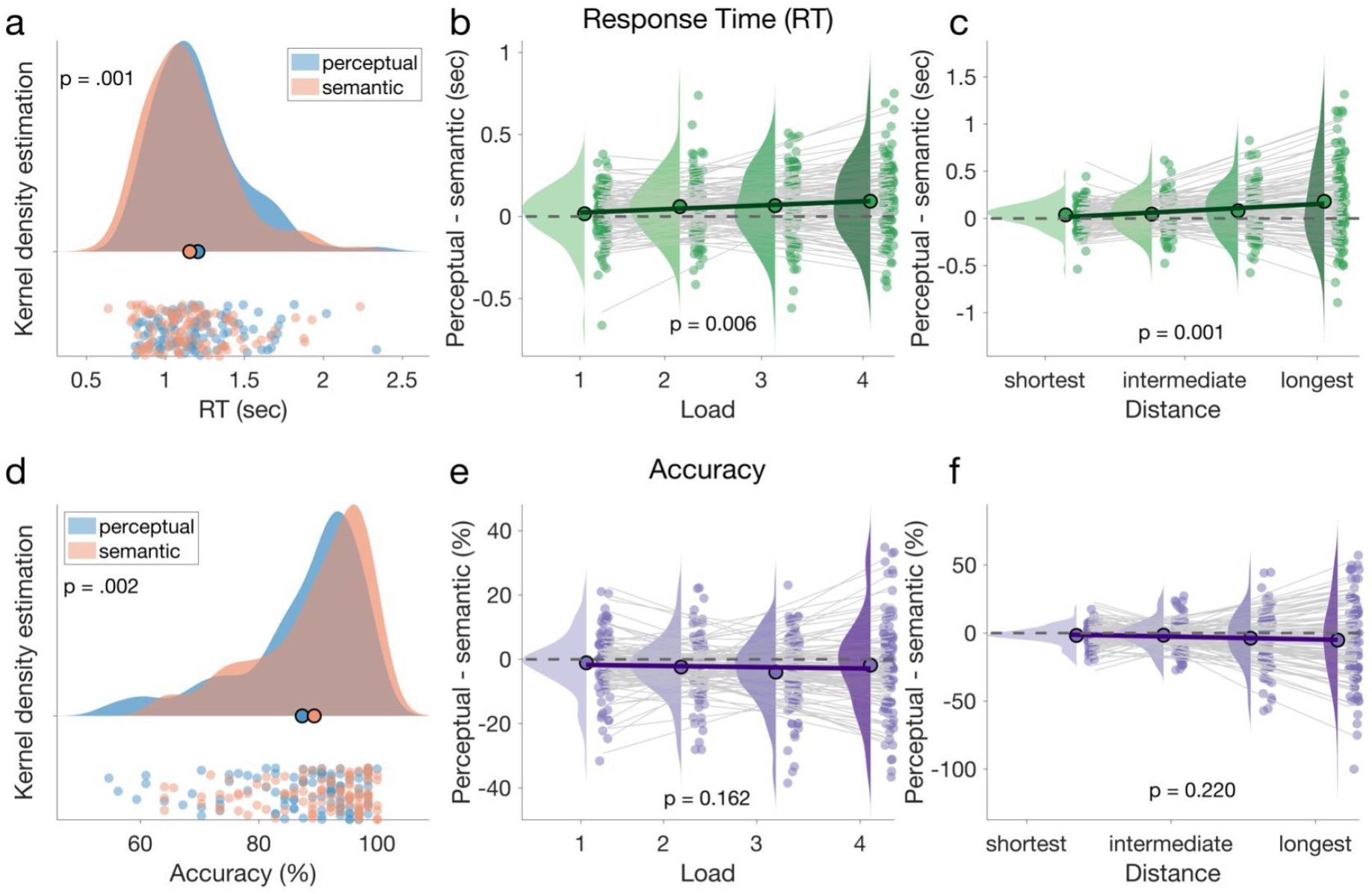
Response times and accuracy results, Experiment 2. Same conventions as *Fig. 2*. ***a***, We again observed overall faster RTs for the semantic (red) compared to the perceptual stimulus characteristics (blue). ***b***, Like in Exp.1, the RT difference (perceptual - semantic) increased with load (here, 1-4 items). ***c***, The RT difference likewise increased with the temporal distance between encoding and test ***d-f***, The accuracy results mirrored those in Exp. 1, with overall higher accuracy of semantic reports (*d*) but no significant effect of load (*e*) or distance (*f*) on the accuracy difference (perceptual-semantic).

Next, we examined the extent to which the prioritization of semantic aspects was modulated by WM load (1-4 items) and/or by the time between item presentation and recall (i.e., the time during which the item was presumably partially unattended). We again observed a significant increase of semantic prioritization in RTs both with load (Fig. 3b; load 1, M = 17ms, SE = 16ms; load 2, M = 60ms, SE = 19ms; load 3, M = 66ms, SE = 21ms ; load 4, M = 93ms, SE = 23ms; Z = 2.74, r = .27, *p* = .006; Wilcoxon signed-rank test of linear slope against zero) and with temporal distance from encoding (Fig. 3c; shortest, M = 38ms, SE = 13ms; second shortest, M = 46ms, SE = 19ms; second longest, M = 82ms, SE = 25ms; longest, M = 182ms, SE = 44ms, Z = 3.43, r = .34, *p* < .001). Like in Exp. 1, the corresponding patterns in accuracy (Fig. 3e and f) were non-significant (load 1, M = -.11%, SE = .87%; load 2, M = -2.39%, SE = .92%; load 3, M = -3.97%, SE = 1.08%; load 4, M = -1.80%, SE = 1.44%, Z = -1.40, r = -.14, *p* = .16; distance: shortest, M = -1.77%, SE = .66%; second shortest, M = -1.56%, SE = 1.08%; second longest, M = -3.80%, SE = 1.90%; longest, M = -5.21%, SE = 2.92%, Z = -1.23 r = -.12, *p* = .22).

In summary, using a more conventional WM task layout with spatial cueing, Exp. 2 replicated the finding of a prioritization of semantic aspects with increasing memory load, which could be related to increasingly unattended storage while additional memory items are being processed. In both experiments, the effects were robustly evident in RTs and were only insignificantly indicated in accuracy. In further analysis, to examine the performance differences between the individual load- and distance conditions with increased power, we combined RT and accuracy into a single performance score.

### Evidence for a categorical difference between single- and multi-item conditions

We used a Balanced Integration Score (BIS)^48^, which quantifies performance in terms of both (shorter) RT and (higher) accuracy with equal weight (see ref.^49^ for an empirical test of BIS compared to other measures). Semantic prioritization was quantified as the difference in BIS for the semantic as compared to the perceptual probe questions. The analysis confirmed that the prioritization of semantic aspects increased with working memory load in both experiments (Fig. 4a,c; Exp. 1, Z = 2.51, r = .29, *p* = .012; Exp. 2, Z = 2.80, r = .28, *p* = .005, Wilcoxon signed-rank tests of linear slopes against zero). A similar pattern was again also evident when we grouped trials according to the temporal distance between encoding and test (Fig. 4b,d). In this latter analysis, however, the linear trend was statistically reliable only in Exp. 2 (Z = 2.93, r = .29, p = .003; Exp. 1: Z = 1.59, r = .18, p = .11).

**Figure 4.**
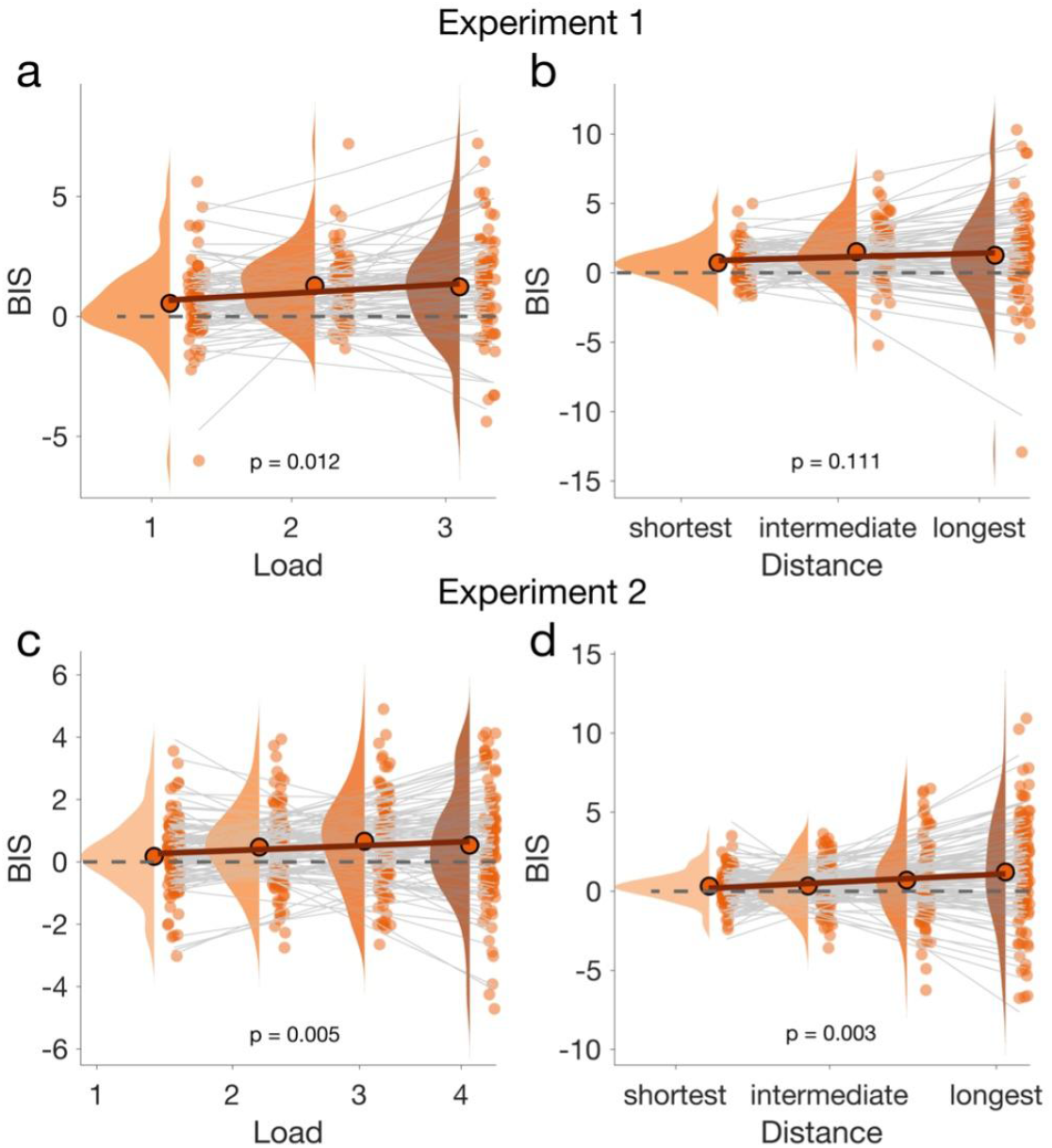
Balanced integration score (BIS) results for Experiments 1 and 2. ***a***, *S*emantic prioritization in terms of BIS difference (semantic-perceptual) across load conditions in Exp. 1. positive values (y-axis) indicate more semantic prioritization. Same conventions as in Figs. 2-3. ***b***, Same as *a*, but with trials grouped according to the distance between encoding and test. The increase in semantic prioritization with distance failed to reach significance. ***c-d***, same as *a-b*, for Exp. 2. Semantic prioritization increased significantly both with load (*c)* and with distance (*d*).

Using BIS, we also explored the possibility that the effects of load were not linear, but rather step-like. Specifically, our results might reflect that accessing a single item in the focus attention (i.e., load 1) is categorically different from retrieving one of multiple items (load>1). Indeed, when repeating the analyses in Fig. 4a,c while excluding the load 1 conditions, the load-effects became non-significant (both Z < .53, both r < .06, both p >.59). As a control, when instead excluding the largest load conditions (load 3 in Exp. 1 and load 4 in Exp. 2), we still observed a robust increase in semantic prioritization across the remaining load levels in both experiments (Exp. 1: Z = 3.88, r = .45, p < .001; Exp. 2: Z = 2.95, r = .29, p = .003). Together, these results suggest a categorical difference in semantic prioritization between single- and multi-item conditions. Lastly, we also examined evidence for or against semantic prioritization in the individual load conditions using Bayesian t-tests against zero^50^ (i.e., no prioritization). Under load 1, we observed only moderate evidence for semantic prioritization in Exp. 1 with verbal cues (BF_10_ = 5.97) and anecdotal evidence for the null-hypotheses (i.e., no prioritization) in Exp. 2 with spatial cues (BF_10_ = .35). Under all higher load conditions (loads >1), in contrast, we observed extreme evidence for semantic prioritization in Exp. 1 (load 2, BF_10_ > 1000; load 3, BF_10_ > 1000), and moderate to extreme evidence for semantic prioritization in Exp. 2 (load 2, BF_10_ = 74.20; load 3, BF_10_ = 763.05; load 4, BF_10_ = 6.83). Together, these results suggest a special status of single-item maintenance as a condition under which semantic prioritization was weak or absent.

To summarize, in both experiments we found clear evidence for a prioritization of semantic over perceptual aspects in accessing information in multi-item WM. Little or no prioritization of either semantic or perceptual aspects was evident in single-item maintenance. Of note, in none of the conditions did we observe faster reporting of the visuo-perceptual than the semantic aspects of the memory items, which clearly distinguishes the accessing of visual working memories from the processing of sensory information in perception.

## Discussion

Higher cognition critically relies on a working memory to keep information available for upcoming tasks. Here, we examined the extent to which the temporal dynamics of accessing information in WM resemble those in visual perception, or those in episodic LTM retrieval, and/or those of selecting action plans. We found response dynamics consistent with selecting prepared actions (see below) when only a single item was to be maintained in WM. However, when multiple items were to be maintained, the response dynamics more closely resembled those in episodic LTM retrieval, where semantic/conceptual aspects are recalled faster than visuo-perceptual details^31,32,42^.

The response dynamics in our multi-item WM tasks were opposite to those commonly observed in visual perception, where low-level visual features are processed faster than high-level conceptual aspects^33–36^. Importantly, the typical time course of visual perception has already been demonstrated in RTs in previous work, using the same response categories (visual: photo/drawing; semantic: animate/inanimate) and stimuli as the current study^31^. Our present finding of semantic prioritization in multi-item WM thus indicates a reversal, or “flip”, of processing dynamics compared to visual perception.

Evidence for a reversal of perceptual processing streams was previously reported in studies of LTM where items were to be recalled only after intervening distractor tasks (which are assumed to prevent WM contributions to recall^51^), and/or after considerable delays. Lifanov et. al^32^, for instance, found increased semantic prioritization 48 hours after studying the memory items (with materials and procedures otherwise similar to our Exp. 1). In contrast, our present results indicate a reversal of processing already within the typical timescales (few seconds) and capacity limits (4 or less items) of WM^22,44^; cf. ref. ^52^ (for related findings in studies of visual imagery, see refs.^37,38^). One possible interpretation is that multi-item WM may recruit similar processes as episodic LTM^8^.

It has previously been hypothesized that a reversal of perceptual processing streams in LTM retrieval might be attributable to the anatomical connections of medial temporal lobe structures critical to episodic memory^31,42^. The proposal is that upon hippocampal pattern completion, memory reconstruction first proceeds from medial temporal regions to nearby multisensory areas which may hold high-level conceptual abstractions, before perceptual details may be reconstructed in anatomically more distant (e.g., early visual) sensory areas. Since we did not record neural activity in the present study, we can only speculate whether such a scenario, which critically involves medial temporal areas, may also explain the semantic prioritization we observed in our multi-item WM tasks.

While contemporary studies of WM in animals and humans have focused mostly on fronto-parietal and sensorimotor cortices (for reviews, see refs.^9,53^, it is worth noting that potential contributions of medial temporal LTM processes have been intensely discussed in the WM literature. For instance, a review by Jeneson & Squire^54^ concluded that medial temporal LTM processes may come into play when WM capacity is exceeded or when attention is temporarily diverted from a memorandum (see also ref.^55^). Similarly, Beukers et al.^8^ proposed that “unattended” or “activity-silent” WM storage^27,56^ may be accomplished through episodic LTM and potentially involve the medial temporal lobes. In the present tasks, all items were equally task-relevant. However, it can be argued that even in our fairly standard multi-item WM tasks (see Experiment 2), individual items had to be temporarily (or partially) unattended, at least while encoding subsequent memory items (i.e., loads >1). As such, the semantic prioritization in our multi-item conditions might likewise be explained through periods of inattention, which would be consistent with a recruitment of LTM-like processes for “unattended WM”.

Although the response dynamics in our multi-item conditions resembled those of episodic LTM retrieval, we exert caution in interpreting the results as direct evidence for a contribution of LTM, as a memory system, to multi-item WM. Using different approaches, two recent WM studies reported evidence for LTM contributions, in terms of a build-up of proactive interference, only for WM loads > 3^57,58^, in ostensible conflict with our present finding of semantic prioritization already at loads > 1. While it remains to be shown what constitutes a definite marker of LTM contributions in behavioral WM tasks, our present findings are open to the alternative interpretation that semantic prioritization is not unique to episodic LTM retrieval, but a more general aspect of cued recall, including from multi-item WM.

We found semantic prioritization to be relatively weak or absent when only a single item was to be maintained (load 1 trials). Our interpretation is that on these trials, participants may have directly accessed the response plans, potentially in a (pre-)verbal format (e.g., “photo” and “animal”), associated with the only item in WM.

In this case, we expected no differences in response times (perceptual vs. semantic), since either of the two response plans could be encoded in the same (e.g., verbal or lexical) format. More specifically, e.g., on load 1 trials in Experiment 2, the WM content may have taken the form of response contingencies (e.g., if probed “animal”, select “yes”; see ref.^43^), which would not qualitatively differ between the perceptual and the semantic probe questions. The load 1 results further indicate that the semantic prioritization observed under higher loads was not due to participants generally focusing more on the items’ semantic aspects (i.e., already during item encoding).

Why would participants not also rely on prepared response plans under higher loads? One possible explanation is that the cognitive demand of maintaining concrete response plans would increase rapidly with the number of items. For instance, under load 3, already six prospective responses (or verbal labels) would need to be maintained and correctly linked (pairwise) to the different cue words or -locations. This would seem an inefficient strategy, also taking into account that the cued test performance in our tasks (e.g., with spatial retrieval cues in Exp. 2) would likely not benefit from maintaining the WM information in serial order (e.g., through rote rehearsal of verbal labels). Under high loads, thus, it seems more efficient to instead rely on an episode-like memory of the presented stimuli.

A central question in WM research is why, in healthy individuals, the phenomenological experience of remembering a stimulus is clearly distinguished from currently perceiving it, despite the “sensory recruitment” of similar brain areas and processes. Our findings may provide a new perspective on this question. In none of the WM conditions did we find the response dynamics observed in visual perception, where low-level sensory aspects are reported before high-level conceptual aspects^31^. We speculate that even if WM contents would be represented in the exact same brain circuits as sensory percepts, they would still give rise to a different phenomenological experience when (re-)constructed in a different (e.g., reverse) temporal order. In other words, the temporal dynamics by which mental representations emerge, on the order of a few hundred milliseconds, might be critical for distinguishing what is seen from what is remembered^42^. While speculative, such a perspective may be useful in studying states of disease or drug effects where the distinction of memories and percepts is blurred, e.g., during hallucinations^59,60^.

In summary, our results add to ongoing discussions about the putative nature of “working” memories, including the extent to which they are retrospective (in terms of a sustained ‘copy’ of past sensory inputs) or prospective (in terms of an emerging action plan or response contingency^43,61^) or both, and how they are distinguished—or not—from long-term memories. Our study provides new evidence for LTM-like dynamics in a typical WM task context (sub-span multi-item maintenance). Distinct, presumedly response-oriented dynamics were observed when a single item was maintained under full attention. Together, the results are consistent with the notion of a single-item focus of attention^30^ within which WM may transform stimulus information into concrete response plans. Information outside the current focus of attention may instead reside in formats more akin to episodic memories, or potentially even recruit episodic memory proper.

## Methods

### Experiment 1

#### Participants

We recruited 109 participants online, of which n=104 completed the experiment (48 female, 56 male; mean age 25.6 years, SD ± 4.8 years). The experiment was terminated prematurely for n=5 participants who failed to pass attention checks (see below). Participation was with informed consent via the Prolific platform (https://www.prolific.ac/). The eligibility criteria were that participants had to be between 18 and 35 years old, fluent in English, normal, or corrected-to-normal vision, and have a minimum approval rate of 95 on Prolific. Participants were reimbursed with £2.5 for completing the experiment which lasted approximately 20 minutes. Participants whose accuracy did not significantly exceed chance level (p<.05, Binomial test against 50% correct responses) in either the semantic or perceptual judgments (n = 29 participants) were excluded. Thus, n=75 participants remained for analysis. The experiment was approved by the Internal Review Board (IRB) of the Max Planck Institute for Human Development.

#### Stimuli

The experimental stimuli comprised 126 pictures (63 depicting animals, 63 depicting inanimate objects). An additional 8 pictures were used for instructions and practice trials. A detailed description of the stimulus set is provided in ref. ^31^. The majority of pictures (96) were from the BOSS database^62^ and the remaining pictures from royalty-free online databases. For each of the original pictures (which are color photographs), a line-drawing version (black-and-white) was created using GNU image manipulation software (http://www.gimp.org). For each participant, half of the animals and half of the inanimate objects were shown as line drawings (randomly assigned) and the others as color photographs. The stimuli thus differed in two orthogonal dimensions, visuo-perceptual (photograph/drawing) and semantic (animate/inanimate; Fig. 1a). In addition, 126 action verbs were used as cue words, which were randomly assigned to the individual pictures for each participant.

#### Task

Each trial started with a central fixation cross (0.5 s), after which 1-3 verb-picture pairs were sequentially presented (Fig. 1b). Each verb was shown for 1s, followed by a picture for 3s. Participants were asked to vividly associate the action verb with the picture (e.g. mentally “sculpting” the lovebird in Fig. 1b). On trials with more than one verb-picture pair (loads >1), the pictures were pseudo-randomly chosen so that they never were all of the same (perceptual or semantic) type. After 1-3 pairs (randomly varied) a response screen appeared (Fig. 1b, *right*) which showed one of the previously presented action verbs (randomly selected) at center, surrounded by four response options (“photo”, “drawing”; “animal”, “no animal”) at the top, bottom, left, and right of the screen. The spatial positions of the options were randomly shuffled on each trial. Participants were asked to remember the picture associated with the action verb, and to select two of the response options (in self-determined order) to report its characteristics (perceptual and semantic) using arrow keys. Upon selection of an option, its font was grayed out and it became inactive. Participants were given 4s time to make their selections. Subsequently, as feedback, the font color of the selected options turned to green (correct) or red (incorrect) for 1.5s until the next trial started. Trials in which participants selected two mutually exclusive options (e.g., “animal” and “no animal”) or failed to respond within the allotted time were excluded from analysis (0.7% of trials on average; min = 0%, max = 4.76%). Note that in Exp 1, each regular trial yielded two responses (one perceptual, one semantic), both of which were included in analysis. We performed additional analyses to examine whether effects were attributable to systematic differences in response order (see *Results*).

#### Procedure

After written instructions, participants performed five practice trials to familiarize themselves with the task. Participants were free to repeat the instructions and practice trials until they felt confident at performing the task. Thereafter, each participant performed 63 experimental trials (21 in each load condition). After the 4th, 10th, and 16th trial, a brief attention check was performed. For this, a large filled triangle was presented centrally in either green, blue, or red color and participants were asked to name the color via button press. If a participant failed on any of the attention checks, the experiment was aborted (see above, *Participants*). After 30 trials, participants could take a short break (self-paced).

### Experiment 2

#### Participants

One hundred and ten participants were recruited online, of which n=109 completed the experiment (48 female, 61 male; mean age 25.9 ± 4.4 years)., with the same eligibility criteria as in Experiment 1. For one participant (n=1), the experiment was terminated prematurely due to failed attention checks (see below). Participants received a reimbursement of £5.5 for completing the experiment which lasted approximately 40 minutes. Participants who failed to perform above chance level (n = 5; same criteria as in Experiment 1) were excluded, leaving n = 103 participants for analysis.

#### Stimuli, Task, and Procedure

The design of Exp. 2 was similar to Exp. 1, with the following differences. The stimulus set was extended with additional pictures from the BOSS database for a total of 154 pictures (77 depicting animals, 77 depicting inanimate objects), which were again presented either as photographs or line-drawings. The line-drawing versions of the additional pictures were created using an online image manipulation tool (https://www.rapidresizer.com). On each trial, 1-4 pictures were sequentially presented (1.5 s/picture) at different locations on the screen (Fig. 1c). The locations were randomly selected from 16 equidistant positions on an (invisible) circle around screen center. On trials with 2 or more pictures, the positions were selected to be at minimum 45° apart from each other. After 1-4 pictures (randomly varied), a probe question appeared at one of the previous stimulus locations (randomly selected), and participants were asked to respond to it via yes/no keys, to indicate the characteristics of the picture remembered at that location. The probe question was any of “animal?”, “object?” (semantic dimension), “photo?”, or “drawing?” (perceptual dimension), and it was randomly varied whether the semantic or the perceptual dimension was probed first. Participants were given 4s time to respond, after which a second probe question (about the other dimension) was presented for 4s. After either response, the probe question’s font turned to green (correct) or red (incorrect) to provide feedback. We only analyzed responses to the first probe question, the dimension of which (perceptual or semantic) could not be anticipated by the participants. Trials in which participants did not respond in time to the first probe question were excluded from analysis (1.2% of trials, min = 0%, max = 22%). Each participant performed 128 trials. On 38 trials, only one picture was presented (load 1 trials), and 30 trials were performed in each of the higher load conditions (2-4; all conditions randomly intermixed). Attention checks and breaks were implemented as in Exp. 1, however, we used a more lenient termination rule and aborted the experiment only when a participant failed on two of the three attention checks.

Both experiments were created using PsychoPy^63^ and custom-written Python and JavaScript code.

## Data Analysis

All analyses were performed in MATLAB 2020a (©The Mathworks, Munich, Germany). To examine in- or decreases in semantic prioritization with load- and distance levels (1-3 in Exp. 1 and 1-4 in Exp. 2), we tested whether the slope of a linear fit (computed on the individual subject level using the function glmfit.m) differed significantly from zero, using Wilcoxon signed-rank tests (two-tailed) on the group level. For consistency and parsimony, we used the same approach also when only two data points were analyzed (see analysis of linear vs. step-like effects in Fig. 4a) instead of performing equivalent pairwise comparisons. Complementary Bayesian t-tests were performed using the bayesFactor toolbox (Krekelberg, 2022; https://github.com/klabhub/bayesFactor) as detailed in *Results*. The BIS was calculated by standardizing (z-scoring) the RTs on correct trials and the accuracies relative to their mean and standard deviation over all conditions, and then subtracting within each condition the RT score from the accuracy score ^48^.

## Data and code availability

All data and code supporting this study are available at https://doi.org/10.5281/zenodo.6593174.

## Author contributions

CK, JLD, and BS designed the experiments. CK implemented and performed the experiments with contributions from JLD. CK and JLD analyzed the data and CK visualized the results. CK, JLD, and BS wrote the paper.

## Acknowledgements

We wish to thank Maria Wimber for stimulating discussions, Thomas Graham for helpful comments on the manuscript, Philip Jakob for technical assistance, and the Center of Adaptive Rationality (Director: Ralph Hertwig) for general support. This research was supported by a European Research Council (ERC) Consolidator Grant ERC-2020-COG-101000972 (BS).

